# *BAPC-assisted* CRISPR/Cas9 System: Targeted Delivery into Adult Ovaries for Heritable Germline Gene Editing (Arthropoda: Hemiptera)

**DOI:** 10.1101/478743

**Authors:** Wayne B. Hunter, Maria T. Gonzalez, John Tomich

**Affiliations:** USDA, ARS, 2001 South Rock Road, Fort Pierce, FL, 34945. USA.; John Tomich, Department of Biochemistry and Molecular Biophysics, Kansas State University, Manhattan, KS, 66506. USA.

**Keywords:** Asian citrus psyllid, Citrus, Gene edit, RNAi Leafhoppers, Huanglongbing, management, pest control

## Abstract

Innovative gene targeting strategies are often limited in application across arthropod species due to problems with successful delivery. In hemipterans, embryonic injections often used to deliver CRISPR components fail due to nearly complete embryo mortality. The Asian citrus psyllid, *Diaphorina citri*, Kuwayama, (Hemiptera: Liviidae), is the vector for a pathogenic bacterium, *Candidatus* Liberibacter asiaticus, CLas, which is devastating the U.S. citrus industries. The disease called, Huanglongbing, HLB, (aka. Citrus greening disease), is transmitted during psyllid feeding. Infection causes severe tree decline, loss of fruits, and eventually tree death. The citrus tree pathogen, CLas, is a fastidious alpha-proteobacterium, which has spread into all citrus growing regions worldwide. The economic losses are estimated in the billions of dollars, in U.S.A., Brazil, and China. Innovative technologies aimed at reducing psyllid populations using targeting RNA suppression, like RNAi, or gene-editing tools, like CRISPR/Cas9 have potential to reduce psyllid vectors and the pathogen in a highly specific manner. Breakthroughs that improve gene editing in psyllids, such as the *BAPC-assisted*-CRISPR/Cas9 System, enabled delivery by injection of CRISPR/Cas9 components directly into nymphs and adult females. Injection near ovaries produced heritable germline gene editing in subsequent generations. This method opens the world of gene editing across arthropods and bypasses the need for microinjection of eggs. Effective development of therapeutic treatments to reduce insect vectors, and stop pathogen transmission would provide sustainable citrus and grapevine industries.

## INTRODUCTION

Biotechnologies provide techniques that greatly improve the level of safety and target specificity in the management of pests and pathogens (Bhaya et al, 2011; Garneau et al, 2010; Fire et al, 1998; Doudna & Charpentier 2014). These techniques include targeted RNA suppression, gene regulation, and gene editing in all organisms: bacteria, plants, animals, and humans. As traditional chemical insecticides fail to provide adequate pest management, due to development of chemical resistance, dependence upon biotech strategies for management have become the best options for development of therapeutic treatments to reduce arthropod vectors, the pathogens, or to cause disruption of vector pathogen acquisition and transmission (Andrade & Hunter 2016; Baum & Roberts 2014; Gantz & Akbari 2018; Hunter & Sinisterra-Hunter 2018; Kolliopoulou et al, 2017; Roberts et al, 2015; Petrick et al, 2013;2016; Scott et al, 2013; Sinisterra-Hunter & Hunter 2018; Taning et al, 2017; Zotti et al, 2018). The rapid emergence of gene editing techniques, like CRISPR/Cas9, Clustered regularly interspaced short palindromic repeats (CRISPR) and the CRISPR-associated protein, Cas9, provide precise editing of genes across all species (Doudna & Charpentier 2014; Peng et al., 2014; Wang et al. 2016). As a natural mechanism in bacteria, numerous studies have described the mode of function of the CRISPR defense system that is an adaptive mechanism, which enables bacteria to suppress invading viruses (Garneau et al, 2010; Doudna & Charpentier 2014). The technology has now been co-adapted to target genes within insect pests (Taning et al, 2017). The relative ease of use, normally by injection into eggs/embryos, combined with improved methods for increase efficacy in CRISPR/Cas9 systems has led CRISPR/Cas to become the primary gene editing tool in the life sciences. For reviews on CRISPR/Cas9 systems see: (Boettcher & McManus 2015; Dominguez et al. 2016; Gupta & Shukla 2016; La Russa & Qi 2015; Liang et al. 2015; Taning et al, 2017; Wang et al. 2016; Wilson & Doudna 2013). Reviews on other gene editing systems with Zinc finger nucleases and TAL effector nucleases, TALENS, see: (Gaj et al, 2013; Bortesi & Fischer 2015; Markert et al, 2016).

CRISPR/Cas systems have demonstrated important applications in agriculture, increasing the options for the management of arthropod pests, insect vectors, and treatments against pathogens of plants, animals and humans (Chen et al, 2016; Chen et al, 2017; Cui et al, 2017; Taning et al, 2017; Sun et al, 2017; Gantz & Akbari 2018; Gundersen-Rindal et al, 2017; Sinisterra-Hunter & Hunter 2018). However, one of the hurdles for rapid adoption in arthropods has been the reliance on embryo injections (Li et al, 2017), which is often unsuccessful in many arthropod species (Bortesi & Fischer 2015; Boettcher & McManus, 2015; Chaverra-Rodriguez et al, 2018; Gregory et al, 2016). Thus, an improved delivery method is needed if gene editing is to be realized for many arthropod species. This is especially true within the Hemiptera, which have few successful demonstrations of embryonic transformation through microinjection of eggs.

In 2018, Hunter described a direct method that delivered and produced gene editing using injections into the abdomens of nymphs, pupae, and adults of hemipterans. Specific examples were the Asian citrus psyllid (*Diaphorina citri*, Kuwayama, Hemiptera: Liviidae) and glassy-winged sharpshooter leafhopper (*Homalodisca vitripennis*, (Germar): Hemiptera: Cicadellidae) (Hunter et al, 2018ab). Previous attempts injecting thousands of eggs failed. Switching to injection of 5^th^ instar nymphs and adult females, produced the first successful trials of gene knockouts in psyllids used the Cas9 protein, co-injected with two sgRNA, producing about 30% surviving G0 and G1 mutants. To improve the system, experiments evaluated the incorporation of BAPC-assisted delivery (Hunter, Gonzalez, Tomich 2018).

### *BAPC-assisted* Delivered RNA interference In Arthropods

The **<u>B</u>ranched <u>A</u>mphiphilic <u>P</u>eptide <u>C</u>apsules (BAPC)**, are a new class of inert, self-assembling peptide nano-capsular spheres (Phoreus Biotechnology, Inc., Olathe, KS, USA). The peptide-based nano-assemblies show promise as nano-delivery vehicles for the safe, targeted transport of drugs, plasmids, dsRNA, and siRNA, to specific tissues and organs with minimal off target accumulation (Gudlur et al, 2012; Sukthankar et al, 2013; 2014; Avila et al, 2015).

Studies on RNAi report that incorporation of BAPC with dsRNA to specific beetle genes, caused significant increase in mortality compared to controls upon ingestion (Avila et al, 2018). In those studies, BAPC-dsRNA was fed to *Tribolium castaneum* (Coleoptera), and the pea aphid, *Acyrthosiphon pisum* (Hemiptera). The authors report an improved delivery of dsRNA into cells most likely due to presence of the BAPC, which prevents degradation by nucleases, producing slower controlled release of the dsRNA upon entering cells, resulting in increased RNAi efficacy and subsequent increased mortality of the insects.

Based on the physical properties of BAPC with nucleic acids we hypothesized that BAPC mixed with guide RNA’s and CRISPR/Cas9 components would result in improved delivery and produce a new method for heritable germline gene editing suitable for injection into adult ovaries of psyllids and other arthropods (Hunter & Sinisterra-Hunter 2018; Hunter, Gonzalez, Tomich 2018; Hunter et al, 2019).

Similar proofs-of-concepts in flies are reported by Chaverra-Rodriguez et al, (2018) in mosquitoes using the P2C peptide mediated transduction of the Cas9 plasmid from the female hemolymph into the developing mosquito oocytes. Their technology, termed “Receptor-Mediated Ovary Transduction of Cargo (ReMOT Control), was shown to work well within the Order: Diptera (Mosquitoes). Another delivery system reported by Wang et al, (2018), in mammals, used microvesicles. Extracellular vesicles, known as *arrestin domain containing protein 1* [ARRDC1]-mediated microvesicles (ARMMs). These ARMMs, could package and deliver intracellularly of a myriad of macromolecules, including the tumor suppressor p53 protein, RNAs, and the genome-editing CRISPR-Cas9/guide RNA complex into mammalian cells.

### *BAPC-assisted-*CRISPR/Cas9 Delivery System: Adult injection near ovaries for heritable germline gene editing [Hemiptera: *Diaphorina citri*)

Attempts to deliver CRISPR/Cas9 into psyllid eggs proved to be difficult and unsuccessful. Injection directly into the 4^th^ and 5^th^ nymphs, pupae, or adults proved to be easier, and more effective (Hunter, Gonzalez, Tomich 2018). Because previous research with BAPC-assisted delivery of dsRNA and plasmids resulted in efficient delivery into insects and animal cells in cultures (Avila et al, 2018; Sukthankar et al, 2014), an experiment was designed to evaluate the incorporation of BAPC to improve delivery of CRISPR/Cas9 components into arthropod ovaries of adult hemipterans (Psyllids and Leafhoppers) for heritable embryonic gene editing.

#### Gene Selection

*Diaphorina citri*, have at least two Thioredoxins, TRX-1, TRX-2, with variants in the mitochondria and cytoplasm. Thioredoxin participates in various redox reactions and catalyzes dithiol-disulfide exchange reactions. Thioredoxin 2, TRX-2, is preferred over thioredoxin 1, as a reducing substrate of peroxiredoxin-1. Thioredoxin is required for female meiosis and early embryonic development. The functions of at least 30 proteins, including enzymes and transcription factors, the regulation of cellular proliferation and the aging process are regulated by TRX (Yoshida et al, 2005). The guide RNA’s and protospacer adjacent motif (PAM) were identified and confirmed using software (Dharmacon, Inc., (Lafayette, CO, 80026). The CRISPR-associated protein 9 (Cas9) was purchased, and protocols were gleaned from publications on CRISPR (Bassett et al, 2013; Kistler, et al, 2015; Larson, et al, 2013; Zhang & Reed 2017; Garczynski et al, 2017). The ACP gene sequence ID: XM_008487100.1, gene, thioredoxin-2-like (LOC103521994) was from the DIACI_2.0 Genome assembly: https://citrusgreening.org/organism/Diaphorina_citri/genome (Saha et al, 2017).

### CRISPR/Cas9 Injections

Injections with CRISPR components, sgRNA’s, and a Cas9 protein, successfully produced knock-out G0 and G1 mutants (Hunter, Gonzalez, Tomich 2018). Subsequent trials incorporated the aid of the *BAPC-assisted*-CRISPR components, successfully produced heritable knockout G2 mutants. Both these methods successfully demonstrated the first psyllid gene knockouts, KO, with CRISPR/Cas9. Designs to *Diaphorina citri*, Asian citrus psyllid, ACP, (Hemiptera: Liviidae) for the knockout used two gRNAs to direct the Cas9 endonuclease to two sites 556 bp apart (Dharmacon, Inc.). The ACP-TRX-2 KO trials injected 30 nymphs (4^th^ and 5^th^ instar), and 20 adult females per treatment (co-injection of two sgRNAs (100 ng/ µL of each, with 200 ng/µL of Cas9 protein, plus BAPC (0.1 ng/ µL) (Drummond Nanoject III, 3-000-207) (Hunter, Gonzalez, Tomich 2018; Hunter et al, 2018; Hunter & Sinisterra 2018). Insects were injected ventrally, off center of midline in the abdomen (FIG. 1). Post injections psyllids were transferred to a citrus seedling to oviposit (~25 cm tall, sweet orange seedlings). A cohort of 6 psyllid adult females were individually analyzed 7 d post treatments using PCR analyses and sequencing. Primers were designed to bracket the gDNA KO sequence of the TRX ORF at 200 to 250 nt beyond the ends in each direction. The remaining cohort of ACP nymphs, which were TRX-KO mutants took 6 to 8 d longer to eclose to adults. The adult psyllids with TRX-KO had significantly shorter lifespans post eclosion living an average 8.5 d, compared to controls injected with buffer, or GFP-plasmid (FIG. 2), which lived an average of 16 d post eclosion (Hunter Gonzalez, Tomich 2018; Hunter & Sinisterra-Hunter 2018; Hunter et al, 2019). (Plasmid resource, Addgene™: pAc5.1B-EGFP was a gift from Elisa Izaurralde, Addgene plasmid #21181)(Karlikow et al, 2016; Legaz et al, 2015). Adult female insects producing eggs (FIG. 3), when treated also produced G2 mutants. Delivery of a GFP-plasmid, when mixed with BAPC, was successfully ingested and expressed in adult psyllids (FIG. 4).

**Figure 1.**
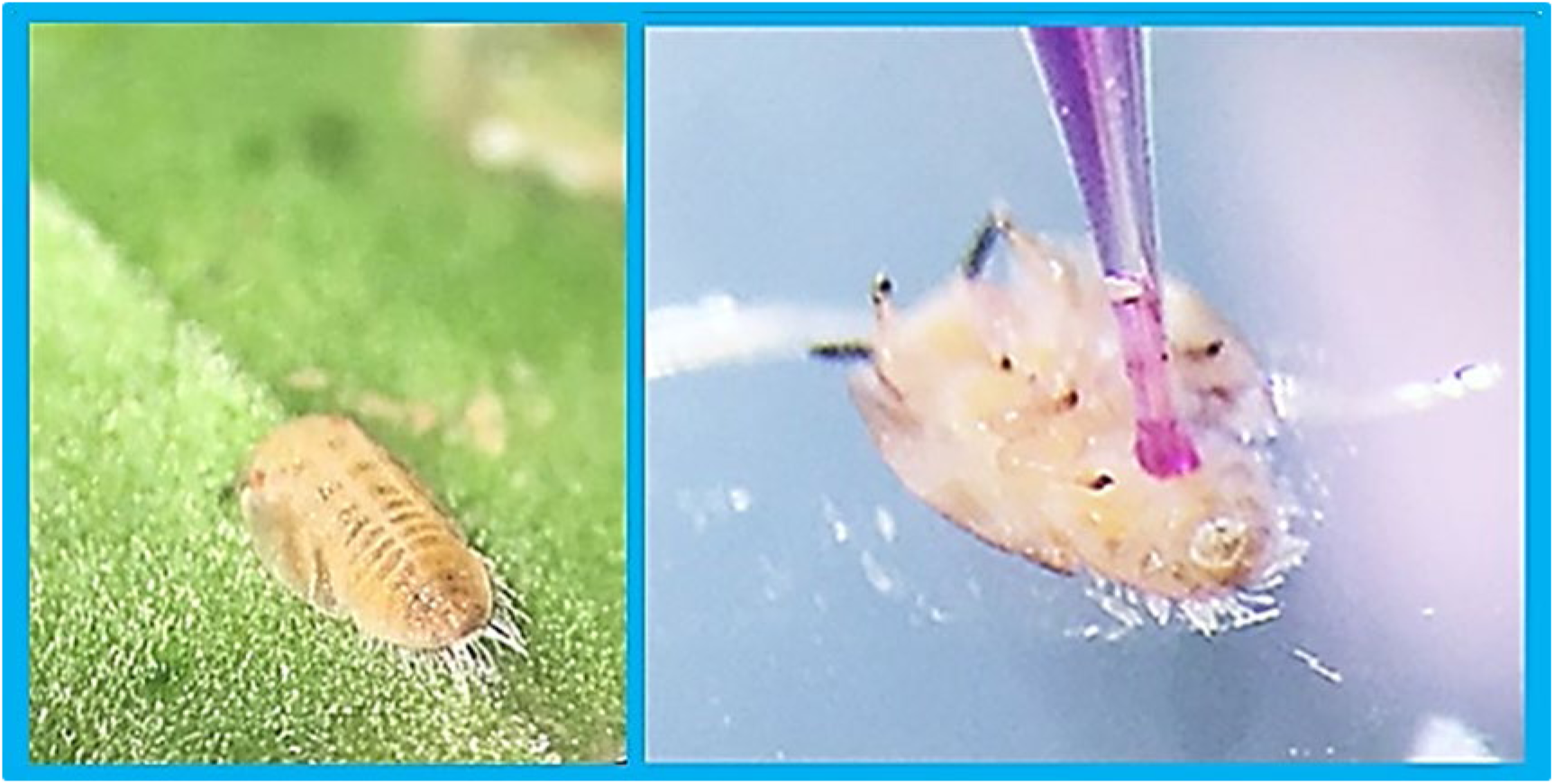
Method of micro-injection of CRISPR/Cas9 components into abdomens of 4^th^, 5^th^ instars and adult female, Asian citrus psyllid, *Diaphorina citri*, (Hemiptera: Liviidae)(Drummond Nanoject III). Nymph on citrus leaf (Left). Nymphs and adults were placed onto solidified, chilled, 1% agar for injections. Shown is artificial dyed solution for easier visualization of method. Abdomen is most proximal, with the head and two dark antennae more distal.

**Figure 2.**
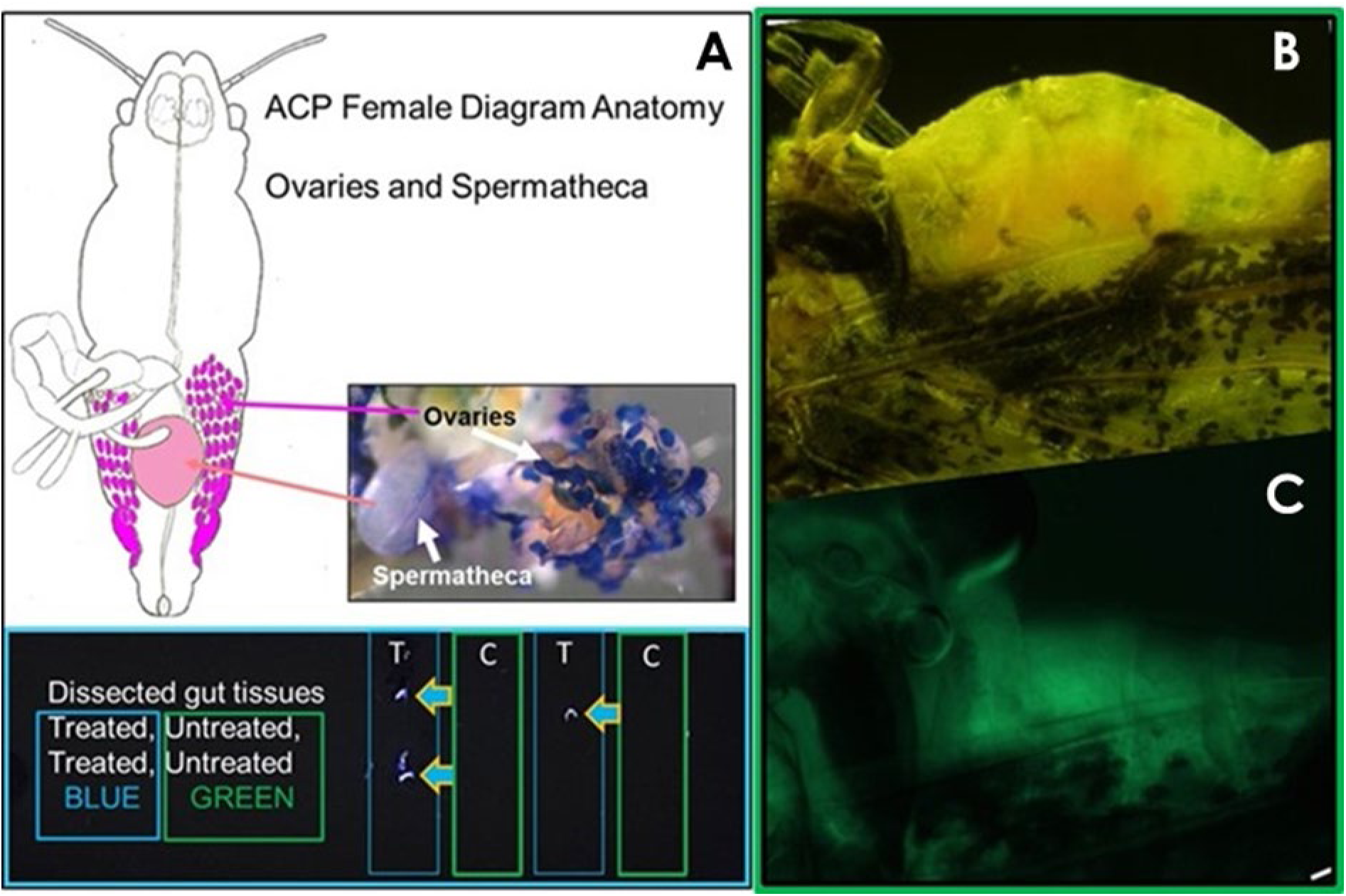
Female adult Asian Citrus Psyllid, *Diaphorina citri* Kuwayama. **A)** Female anatomy showing ovaries, and spermatheca in diagram. Fresh dissection (stained dark blue, Tryptophan). Injection of plasmid-GFP expression, psyllid dissected 8 d post injection. Illumination of Green fluorescent protein, GFP with UV light (Blue Arrows). **B)** The GFP-plasmid (MTG-Dc-1, actin) was injected, **C)** Expression under Dc-Actin promoter identified from *DIACI_2.0 genome,* OGS_0.2v (Saha, et al, 2017). (Plasmid, Addgene, Karlikow et al, 2016).

**Figure 3.**
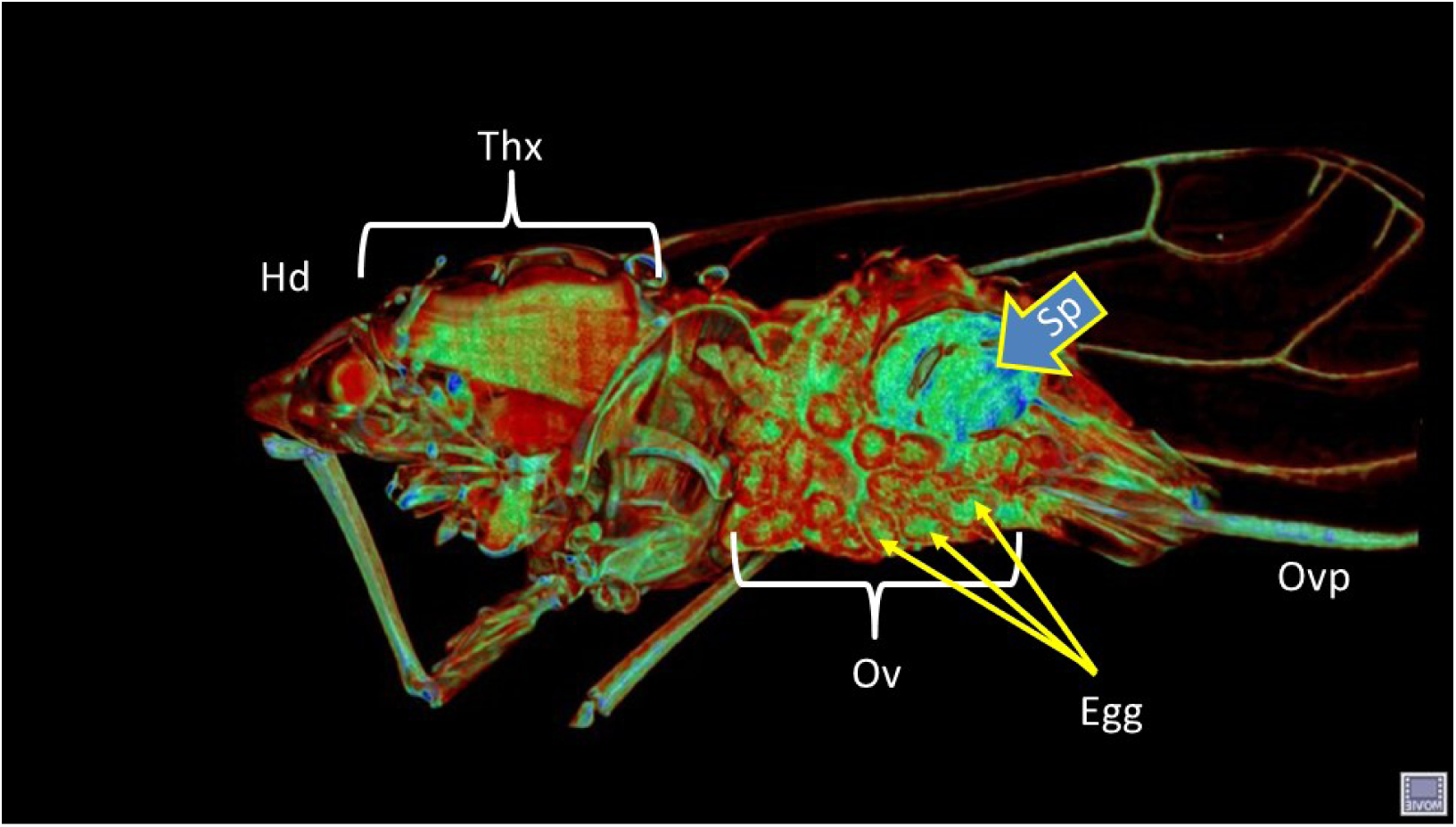
Micro-CT imaging of adult female Asian citrus psyllid, *Diaphorina citri*, Kuwayama (Hemiptera: Liviidae). (Alba-Tercedor, J. and Hunter, W.B. 2016). www.citrusgreening.org [Alba-Tercedor et al, 2018].

**Figure 4.**
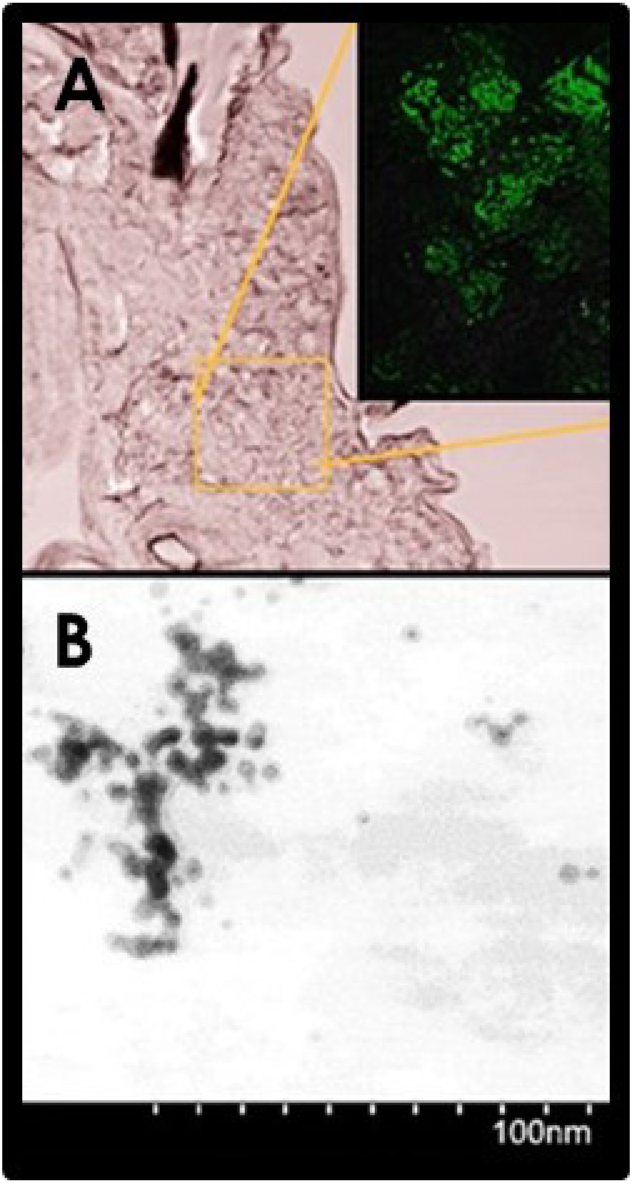
BAPC-assisted delivery of GFP-plasmid post ingestion. Trial using BAPC-labeled with fluorescent Probe Atto488. Oral delivery to adult psyllid, 5 d post feeding, from sucrose solution (25%) plus BAPC in feeding satchet. Thick-section of single psyllid abdomen tissues fixed and parafilm embedded, thick section, (GFP insert, green). Demonstrated delivery and location post ingestion of BAPC in psyllid tissues. Psyllid actin promoter designed as previously mentioned in methods (MTG-Dc-1).

## Conclusions

Advances in biotechnologies, like CRISPR and RNAi provides new sustainable and environmentally friendly strategies to reduce insect vectors, like psyllids (Andrade & Hunter 2016; 2017; Hunter & Sinisterra-Hunter 2018; Taning et al, 2016; Ghosh et al, 2018). These advances will also aid in the management of many other insect vectors and pests (Chaverra-Rodriguez et al, 2018; Darrington et al, 2017; Dong, et al, 2015; 2018; Gantz & Akbari 2018; Ghosh et al, 2018; Kolliopoulou et al, 2017; Sinisterra-Hunter & Hunter 2018; Taning et al, 2017; Zotti et al, 2018).

CRISPR/Cas9 gene editing in hemipterans was shown to be feasible using nymphs and adult psyllids, as the recipient for gene editing CRISPR components. The method was a significant improvement over efforts using injection of eggs. Furthermore, incorporation of *BAPC-assisted* delivery of CRISPR/Cas9 components into nymph and adults provides an innovative breakthrough in hemipteran gene editing. Production of G2 mutants, from *BAPC-assisted*-CRISPR/Cas9 injected, adult female psyllids further supports the viability of this method. Improvements in efficacy, by adjusting component concentration ratios still need to be evaluated across several hemipteran species.

## Acknowledgements

We thank Dr. Steve Garczynski for consultations on CRISPR/Cas9; Ken Sims, Biological Science Technician, sample fixation, paraffin embedding, and Jennifer Wildonger, Technician, sample sectioning, confocal, USDA, ARS, Ft. Pierce, FL., Dr. Michael Boyle, confocal training, Smithsonian Marine Station, Ft. Pierce, FL, and Chris S. Holland, confocal imaging, bioassay trials, ORISE program DOE/USDA. *This research was supported in part by an appointment to the Agricultural Research Service (ARS) Research Participation Program administered by the Oak Ridge Institute for Science and Education (ORISE) through an interagency agreement between the U.S. Department of Energy (DOE) and the U.S. Department of Agriculture (USDA). ORISE is managed by ORAU under DOE contract number DE-SC0014664. All opinions expressed in this paper are the author’s and do not necessarily reflect the policies and views of USDA, ARS, DOE, or ORAU/ORISE.* USDA, ARS 2015-2018. Funding in part from: National Institute of Food and Agriculture, USDA, Specialty Crops Research Initiative/Citrus Disease Research & Extension. Award #2015-70016-23028, Developing an Infrastructure and Product Test Pipeline to Deliver Novel Therapies for Citrus Greening Disease, Lead Dr. Susan Brown. Kansas State Univ., Manhattan, KS, & USDA, ARS, Fort Pierce, FL.

## Disclaimer

*This article is a US Government work and is in the public domain in the USA. Mention of a trademark or proprietary product is for identification only and does not imply a guarantee or warranty of the product by the US Department of Agriculture. The US Department of Agriculture prohibits discrimination in all its programs and activities on the basis of race, color, national origin, gender, religion, age, disability, political beliefs, sexual orientation and marital or family status*.

## References

Alba-Tercedor, J., Hunter, W.B. 2016. Micro-CT imaging of adult female Asian citrus psyllid, Diaphorina citri, Kuwayama (Hemiptera: Liviidae). www.citrusgreening.org

Alba-Alejandre, I., Hunter, W.B., Alba-Tercedor, J. 2018. Micro-CT study of male genitalia and reproductive system of the Asian citrus psyllid, *Diaphorina citri Kuwayama, 1908* (Insecta: Hemiptera, Liviidae). PLoS ONE 13(8): e0202234. https://Doi.org/10.1371/journal.pone.0202234.

Alba-Tercedor, J., Hunter, W.B., Cicero, J.M.., Sáinz-Bariáin, M., Brown, S.J. 2017. Use of micro-CT to elucidate details of the anatomy and feeding of the Asian citrus psyllid, Diaphorina citri Kuwayama, 1908 (Insecta: Hemiptera, Liviidae).

Micro-CT User Meeting 2017. Pp. 270–285. Royal Belgian Institute of Natural Science, Brussels.

Andrade, C.E., Hunter, W.B. 2016. RNA interference —Natural gene-based technology for highly specific pest control (HiSPeC) In RNA Interference, ed I. Y. Abdurakhmonov (Croatia: InTech), 391–409. http://www.intechopen.com/books/rna-interference/rna-interference-natural-gene-based-technology-for-highly-specific-pest-control-hispec-

Andrade, EA., Hunter, W.B. 2017. RNAi feeding bioassay: development of a non-transgenic approach to control Asian citrus psyllid and other hemipterans. Entomol. Exper. et Applic. 162:389–396. Doi:10.1111/eea.12544

Avila, L.A., Aps, L. R. M. M., Sukthankar, P., Ploscariu, N., Gudlur, G., Šimo, L. Szoszkiewicz, R., Park, Y., Lee, S. W., Iwamoto, T., Ferreira, L., Tomich, J. M. 2015. Alternate supramolecular structures for different ratios of branched amphiphilic cationic peptide /DNA assemblies: Correlation with gene delivery Mol. Pharm. 12(3): 706–715.

Avila, L.A., Chandrasekar, R., Wilkinson, K., Balthazor, J., Herman, M., Bechard, J., Brown, S., Schoenek, B., Dhar, S., Tomich, J.M. Reeck, G.R. 2018. Delivery of lethal dsRNAs in insect diets by branched amphiphilic peptide capsules. J. Contrl Release. 273: 139–146.

Bassett, A.R., Tibbit, C., Ponting, C.P., Liu, J. 2013. Mutagenesis and homologous recombination in *Drosophila* cell lines using CRISPR/Cas9. Biol. Open 3: 42–49.

Bhaya, D., Davison, M., Barrangou, R. 2011. CRISPR-Cas systems in bacteria and archaea: versatile small RNAs for adaptive defense and regulation. Annu. Rev. Genet. 45: 273–297. Doi:10.1146/annurev-genet-110410-132430.

Boettcher, M.; McManus, M.T. 2015. Choosing the right tool for the job: RNAi, TALEN, or CRISPR. Mol. Cell 58:575–585. Doi.org/10.1016/j.molcel.2015.04.028.

Bortesi, L., Fischer, R. 2015. The CRISPR/Cas9 system for plant genome editing and beyond. Biotechnol Adv. 33(1):41–52. Doi:10.1016/j.biotechadv.2014.12.006.

Chaverra-Rodriguez, D., Macias, V.M., Hughes, G.L., Pujhari, S., Suzuki, Y., Peterson, D.R., Kim, D., McKeand, S., Rasgon, J.L. 2018. Targeted delivery of CRISPR-Cas9 ribonucleoprotein into arthropod ovaries for heritable germline gene editing. Nature Communications 9:3008. Doi:10.1038/s41467-018-05425-9.

Chen, L., Wang, G., Zhu, Y. N., Xiang, H., Wang, W. 2016. Advances and perspectives in the application of CRISPR/Cas9 in insects. Zool. Res. 37, 220–228. Doi.10.13918/j.issn.2095-8137.2016.4.220

Chen, S., Hou, C., Bi, H., Wang, Y., Xu, J., Li, M., et al. 2017. Transgenic CRISPR/ Cas9-mediated viral gene targeting for antiviral therapy of Bombyx mori nucleopolyhedrovirus (BmNPV). J. Virol. 91,e02465–e02516. Doi:10.1128/JVI.02465-16.

Cui, Y.B., Sun, J.L., Yu, L.L. 2017. Application of the CRISPR gene-editing technique in insect functional genome studies-a review. Entomol. Exp. Appl. 162:124–132. Doi:10.1111/eea.12530.

Darrington, M., Dalmay, T., Morrison, N.I., Chapman, T. 2017. Implementing the sterile insect technique with RNA interference - a review. Entomologia Experimentalis et Applicata 164:155–175. Doi.10.1111/eea.12575.

Dominguez, A.A., Lim, W.A., Qi, L.S. 2016. Beyond editing: repurposing CRISPR-Cas9 for precision genome regulation and interrogation. Nat. Rev. Mol. Cell Biol. 17:5–15. Doi.10.1038/nrm.2015.2.

Dong, S., Lin, J., Held, N.L., Clem, R.J., Passarelli, A.L., Franz, A.W.E. 2015. Heritable CRISPR/Cas9-mediated genome editing in the yellow fever mosquito, *Aedes aegypti*. PLoS ONE 10:1–13. Doi.org/10.1371/journal.pone.0122353

Doudna, J.A., Charpentier, E. 2014. The new frontier of genome engineering with CRISPR-Cas9. Science 346(6213):1258096. Doi:10.1126/science.1258096.

Fire, A., Xu, S., Montgomery, M.K., Kostas, S.A., Driver, S.E., Mello, C.C., 1998. Potent and specific genetic interference by double-stranded RNA in *Caenorhabditis elegans*. Nature. 391:806–811. Doi:10.1038/35888.

Gaj, T., Gersbach, C.A., Barbas, C.F, III. 2013. ZFN, TALEN, and CRISPR/Cas-based methods for genome engineering. Trends in Biotechnology 31:397–405. doi.org/10.1016/j.tibtech.2013.04.004.

Gantz, V.M., Akbari, O.S. 2018. Gene editing technologies and applications for insects. Current Opinion in Insect Science 28:66–72. Doi.org/10.1016/j.cois.2018.05.006.

Garczynski, S.F., Martin, J.A., Griset, M., Willett, L.S., Cooper, W.R., Swisher, K.D., Unruh, T.R. 2017. CRISPR/Cas9 Editing of the codling moth (Lepidoptera: Tortricidae) *CpomOR1* gene affects egg production and viability. Journal of Economic Entomology, 110(4), 2017, 1847–1855. Doi:10.1093/jee/tox166.

Garneau, J.E., Dupuis, M., Viooion, M., Romero, D., Barrangou, R., Boyaval, P., Fremaux, C., Horvath, P., Magadan, A., Moineau, S. 2010. The CRISPR/Cas bacterial immune system cleaves bacteriophage and plasmid DNA. Nature 468: 67–71.

Ghosh, S.K., Hunter, W.B., Park, A.L., Gundersen-Rindal, D.E. 2017. Double strand RNA delivery system for plant-sap-feeding insects. PLOS ONE 12(2): e0171861. Doi:10.1371/journal.pone.0171861.

Ghosh, S.K., Gundersen-Rindal, D.E., Park, A.L., Hunter, W.B. 2018. Double-stranded RNA oral delivery methods to induce RNA interference in phloem and plant-sap-feeding hemipteran insects. J. Visualized Experiments 135, e57390. Doi:10.3791/57390.

Gregory, M., Alphey, L., Morrison, N.I., Shimeld, S.M. 2016. Insect transformation with piggyBac: getting the number of injections just right. Insect Mol Biol. 25:259–271. Doi:10.1111/imb.12220.

Gudlur, S., Sukthankar, P., Gao, J. Avila, L.A., Hiromasa, Y., Chen, J., Iwamoto, T., Tomich, J.M. (2012) Peptide Nanovesicles Formed by the Self-Assembly of Branched Amphiphilic Peptides. PLOS ONE. 7(9) e45374. Doi:10.1371/journal.pone.0045374.

Gundersen-Rindal, D., Adrianos, S., Allen, M., Becnel, J.J., Chen, Y.P., et al. (28). 2017. Arthropod genomics research in the United States Dept. Agriculture-Agricultural Res. Service: Applications of RNA interference and CRISPR gene-editing technologies in pest control. Trends in Entomology 13: 109–137.

Gupta, S.K., Shukla, P. 2016. Gene editing for cell engineering: trends and applications, Critical Reviews in Biotechnology, Doi.10.1080/07388551.2016.1214557

Hunter, W.B., Clarke, S.V., Sandoval Mojica, A.F., Paris, T., Metz, J.L., Holland, C.S., et al. (9). 2019. Advances in RNA suppression of the Asian Citrus Psyllid Vector of bacteria causing Huanglongbing (Hemipteran vector / Liberibacter pathogens). Chpt. 19. HLB. Book, (eds) Phil Stansly and Jawwad Qureshi. Commonwealth Agricultural Bureau International (CABI Press). (in press).

Hunter, W.B., Metz, J.L., Temeyer, K.B., de Leon, A.P., Pelz-Stelinski, K. Sandoval Mojica, A.F., McCollum, G., Aishwarya, V. 2018a. FANA_ASO Reduces Pathogens and Pest of Fruit Crops: Citrus and Grapevine. International Plant & Animal Genome XXVI Conf., San Diego, CA. Jan. 13-18. USA.

Hunter, WB., Mojica, AS., Paris, T., Miles, G., Metz, J., McCollum, G., Boyle, M., Altman, S., Aishwarya, V, Quereshi, J. 2018b. Antisense Oligonucleotides, F-ASO, and PPMO, new tools to reduce pests and pathogens in citrus and other agricultural crops. 92^nd^ Ann. Meeting Southeastern Branch, Entomol. Soc. America, March 5-7.Orlando, FL. USA.

Hunter, W.B., Gonzalez, M.T., Tomich, J.M. 2018. BAPC-CRISPR/Cas9 System for Heritable Embryonic Gene Editing: Gene Knock-Out, Asian Citrus Psyllid. Proc. The 101st Ann. Meeting Int’l Florida Entomological Society, St. Augustine, FL, USA. July 22-25. http://www.flaentsoc.org/dl/18_HUNTER_BAPC-CRISPR-Cas9_FES%2022July%20.pdf

Hunter, W.B., Sinisterra-Hunter, X. 2018. Emerging RNA Suppression Technologies to Protect Citrus Trees from Citrus Greening Disease Bacteria. Advances Insect Physiology 55:163–199. Doi.org/10.1016/bs.aiip.2018.08.001

Karlikow, M., Goic, B., Mongelli, V., Salles, A., Schmitt, C., Bonne, I., Zurzolo, C., Saleh, M.C. Drosophila cells use nanotube-like structures to transfer dsRNA and RNAi machinery between cells. 2016. Sci. Rep. 6:27085. Doi:10.1038/srep27085.

Kistler, K.E., Vosshall, L.B. Matthews, B.J. 2015. Genome engineering with CRISPR-Cas9 in the mosquito Aedes aegypti. Cell Rep. 11:51–60. https://doi.org/10.1016/j.celrep.2015.03.009.

Kolliopoulou, A., Taning, C.N.T., Smagghe, G., Swevers, L. 2017. Viral delivery of dsRNA for control of insect agricultural pests and vectors of human disease: Prospects and challenges. Front. Physiol. 8:399. Doi:10.3389/fphys.2017.00399.

La Russa, M.F., Qi, L.S. 2015. The new state of the art: Cas9 for gene activation and repression. Mol. Cell. Biol. 35:3800–3809. Doi.10.1128/MCB.00512-15.

Larson, M.H., Gilbert, L.A., Wang, X., Lim, W.A., Weissman, J.S., Qi, L.S. 2013. CRISPR interference (CRISPRi) for sequence-specific control of gene expression. Nature Protocols 8(11):2180–2196. Doi:10.1038/nprot.2013.132.

Legaz, S., Exposito, J.Y., Borel, A., Candusso, M.P., Megy, S., Montserret, R., Lahaye, V., Terzian, C., Verrier, B. A purified truncated form of yeast Gal4 expressed in Escherichia coli and used to functionalize poly(lactic acid) nanoparticle surface is transcriptionally active in cellulo. 2015. Protein Expr Purif. 113:94–101. Doi:10.1016/j.pep.2015.05.009.

Li, M., Au, L.Y.C., Douglah, D., Chong, A., White, B.J., Ferree, P.M., Akbari, O.S. 2017. Generation of heritable germline mutations in the jewel wasp Nasonia vitripennis using CRISPR/Cas9. Scientific Reports 7:901. Doi:10.1038/s41598-017-00990-3.

Liang, X., Potter, J., Kumar, S., Zou, Y., Quintanilla, R., et al. 2015. Rapid and highly efficient mammalian cell engineering via Cas9 protein transfection. J. Biotechnol. 208:44–53. Doi:10.1016/j.jbiotec.2015.04.024.

Markert, M.J., Zhang, Y., Enuameh, M.S., Reppert, S.M., Wolfe, S.A., Merlin, C. 2016. Genomic access to monarch migration using TALEN and CRISPR/Cas9-mediated targeted mutagenesis. G3: Genes, Genomes, Genetics 6: 905–915.

Peng, Y., Clark, K.J., Campbell, J.M., Panetta, M.R., Guo, Y., Ekker, S.C. 2014. Making designer mutants in model organisms. Development 141: 4042–4054.

Petrick, J.S., Brower-Toland, B., Jackson, A.L., Kier, L.D. 2013. Safety assessment of food and feed from biotechnology-derived crops employing RNA-mediated gene regulation to achieve desired traits: a scientific review. Regulatory Toxicol. Pharmacol. 66:167–176. Doi:10.1016/j.yrtph.2013.03.008.

Petrick, J.S., Frierdich, G.E., Carleton, S.M., Kessenich, C.R., Silvanovich, A., Zhang, Y., Koch, M.S., 2016. Corn rootworm-active RNA DvSnf7: repeat dose oral toxicology assessment in support of human and mammalian safety. Reg. Tox. Pharm. 81:57e68. http://dx.doi.org/10.1016/j.yrtph.2016.07.009.

Roberts, A.F., Devos, Y., Lemgo, G.N.Y., Zhou, X. 2015. Biosafety research for non-target organism risk assessment of RNAi-based GE plants. Front. Plant Sci. 6:958. Doi:10.3389/fpls.2015.00958.

Saha, S., Hosmani, P.S., Villalobos-Ayala, K., Miller, S., et al. (44). 2017. Improved annotation of the insect vector of Citrus greening disease: Biocuration by a diverse genomics community. Database 2017:bax032. https://doi.org/10.1093/database/bax032

Scott, J.G., Michel, K., Bartholomay, L., Siegfried, B.D., Hunter, W.B., Smagghe, G., Zhu, K.Y., Douglas, A.E. 2013. Towards the elements of successful insect RNAi. Journal of Insect Physiology 59:1212–1221. http://dx.doi.org/10.1016/j.jinsphys.2013.08.014

Sinisterra-Hunter, X., Hunter, W.B. 2018. Towards a holistic integrated pest management: Lessons learned from plant-insect mechanisms in the field. *Chpt. 10*. Pp.204–226. *In* The Biology of Plant-Insect Interactions: A Compendium for the Plant Biotechnologist. 236 Pgs. Chandrakanth Emani (ed). ISBN 9781498709736-CAT#K25008. CRC Press.

Sukthankar, P., Avila, L.A., Whitaker, S.K., Iwamoto, T., Morgenstern, A., Apostolidis, C., Liu, K., Hanzlik, R.P. Dadachova, E., Tomich, J.M. 2014. Branched Amphiphilic Peptide Capsules: Cellular uptake and retention of encapsulated solutes. Biochim, Biophys. Acta, Biomembranes 1838: 2296–2305.

Sukthankar, P., Gudlur, S., Avila, L.A., Whitaker, S., Katz, B.B., Hiromasa, Y., Gao, J., Thapa, P., Moore, D., Iwamoto, T., Tomich, J.M. 2013. Branched oligopeptides form nano-capsules with lipid vesicle characteristics. Langmuir 29:14648–14654.

Taning, C.N.T., Christiaens, O., Berkvens, N., Casteels, H., Maes, M., Smagghe, G. 2016. Oral RNAi to control *Drosophila suzukii*: laboratory testing against larval and adult stages. J. Pest Sci. 89, 803–814. Doi:10.1007/s10340-016-0736-9

Taning, C.N.T., van Eynde, B., Yu, N., Ma, S., Smagghe, G. 2017. CRISPR/Cas9 in insects: applications, best practices and biosafety concerns. J. Insect Physiol. 98:245–257. Doi.org/10.1016/j.jinsphys.2017.01.007.

Wang, H., Russa, M.L., Qi, L.S. 2016. CRISPR/Cas9 in genome editing and beyond. Annu. Rev. Biochem. 85:227–264. Doi:10.1146/annurev-biochem-060815-014607.

Wang, Q., Yu, J., Kadungure, T., Beyene, J., Zhang, H., Lu, Q. 2018. ARMMs as a versatile platform for intracellular delivery of macromolecules. Nature Communications 9–960. Doi:10.1038/s41467-018-03390-x

Wilson, R.C., Doudna, J.A. 2013. Molecular mechanisms of RNA interference. Annu. Rev. Biophys. 42: 217–239. Doi:10.1146/annurev-biophys-083012-130404.

Yoshida, T., Nakamura, H., Masutani, H., Yodoi, J. 2005. The involvement of thioredoxin and thioredoxin binding protein-2 on cellular proliferation and aging process. Annals New York Acad. Sciences. 1055:1–12. Doi:10.1196/annals.1323.002.

Zhang, L., Reed, R.D. 2017. A practical guide to CRISPR/Cas9 genome editing in Lepidoptera. bioRxiv preprint Apr. 24. Doi.org/10.1101/130344.

Zotti, M., Santos, E.A., Cagliari, D., Christiaens, O., Taning, C.N.T., Smagghe, G. 2018. RNA interference technology in crop protection against arthropod pests, pathogens and nematodes. Pest Manag. Sci. Doi:10.1002/ps.4813.

